# Optimized splitting of RNA sequencing data by species

**DOI:** 10.1101/2021.06.09.447735

**Authors:** Xuan Song, Hai Yun Gao, Karl Herrup, Ronald P. Hart

## Abstract

Gene expression studies using chimeric xenograft transplants or co-culture systems have proven to be valuable to uncover cellular dynamics and interactions during development or in disease models. However, the mRNA sequence similarities among species presents a challenge for accurate transcript quantification. To identify optimal strategies for analyzing mixed-species RNA sequencing data, we evaluate both alignment-dependent and alignment-independent methods. Alignment of reads to a pooled reference index is effective, particularly if optimal alignments are used to classify sequencing reads by species, which are re-aligned with individual genomes, generating >97% accuracy across a range of species ratios. Alignment-independent methods, such as Convolutional Neural Networks, which extract the conserved patterns of sequences from two species, classify RNA sequencing reads with over 85% accuracy. Importantly, both methods perform well with different ratios of human and mouse reads. Our evaluation identifies valuable and effective strategies to dissect species composition of RNA sequencing data from mixed populations.

## 1. Introduction

*Mus musculus* has long been used as a model organism for the study of human biology.^1, 2^ The conserved similarity between human and mouse at genetic, molecular, and physiological levels makes it a powerful tool which effectively contributes to human studies.^3-5^ However, not only do mice respond to drugs in ways that markedly differ from humans,^6^ but mouse strains carrying human genetic variance also frequently fail to fully mimic human phenotypes.^7-9^ Strikingly, a study that examined gene expression in human and mouse tissues reported transcript diversity between comparable tissues from human and mouse,^10^ which indicates fundamental differences between these two organisms. Finally, genome-wide association studies (GWAS) frequently identify disease-associated SNPs in noncoding genome, which may affect regulatory elements,^11^ but noncoding sequence is less conserved between species.^12, 13^ While the mouse remains a valuable model organism for studies of human biology, certain disease-associated phenotypes are better captured and examined in a human genetic background.

Human cells and tissue may be available post-mortem, however, these cannot be used to observe complex biological activities for mechanistic studies.^14^ Particularly for the study of brain, this limitation severely restrains the exploration of complex neuronal network activity, brain development and related neurodegenerative disorders. Studies have reported human-mouse chimeric models with human cells transplanted into mouse.^15-19^ This transplantation system allows us to more closely evaluate donor cellular dynamics *in vivo*.^20, 21^ Similarly, neurons differentiated from human induced pluripotent stem cells (iPSC) are often co-cultured with mouse glial cells for metabolic and trophic support^22-24^ as mouse glia enhance maturation of human induced neurons.^25^ Gene expression studies from either xenograft transplants or co-cultures include sequences from multiple species that need to be resolved.

Bulk RNA sequencing (RNAseq) cannot provide unambiguous species resolution from a mixed population.^26, 27^ Thus, species-specific decomposition is needed after sequencing but before analysis. Single-cell sequencing technologies provide detailed information about the heterogeneity of the cellular mixture but this technique remains expensive, less sensitive, and noisier compared with traditional bulk RNAseq.^28^ Moreover, despite the rapid growth of single-cell level sequencing, the majority of available public databases are based on bulk RNAseq.

Several strategies have been used for processing mixed-species RNAseq data. Regression-based methods decompose expression data from a mixed cell population, but these depend on an estimated reference gene expression profile from purified cell populations using cell sorting.^29, 30^ Other methods require a pre-captured reference profile from single-cell expression data to estimate the composition of cell mixtures,^31-33^ which is not always feasible.^34^ Here we compare methods which directly classify bulk RNAseq reads into human or mouse. The goal is to perform bulk RNAseq using a mixed cell population of both human and mouse and distinguish sequencing reads of each species without a pre-captured reference profile or single-cell sequencing. Our alignment-based approach maps raw sequencing data to pooled reference genomes and compares strategies for choosing the best alignment to classify reads based on species. An alignment-free approach employs a one-dimensional convolutional neuronal network which is commonly used in image classification and text matching problems.^35, 36^ We demonstrate the performance using different sets of mixed RNAseq data of pre-acquired purified human and mouse cells. We found the alignment-free method is accurate when assessing a mixed population with relatively equal proportions of human and mouse cells. This method is time efficient and does not significantly increase time for data processing before analysis. However, alignment-based methods outperform the alignment-free approach, particularly when using the “primary alignment” flag in the initial SAM/BAM files.

## 2. Methods

Our goal was to model RNAseq data from mixed cultures of human neurons induced from iPSC lines co-cultured with mouse glia, since this is a system used in our research.^22, 23, 37^ Therefore, we obtained public FASTQ files of human control interneuron RNAseq data (GEO accession GSE118313, sample GSM3324649, 9,506,181 raw reads)^38^ and mouse astrocytes derived from dorsal root ganglia (GEO accession GSE133745, sample GSM3926526, 80,822,800 raw reads)^39^. Both datasets were paired-end with 75 nt/end.

For alignment-based methods, standard open-source software was used, including HISAT2, Kallisto, Samtools, and Python for scripting.

- HISAT2 v2.2.0 http://daehwankimlab.github.io/hisat2/
- Kallisto v0.46.0 https://pachterlab.github.io/kallisto/
- Samtools v1.3.1 http://www.htslib.org/
- Python v3.7.6 https://www.python.org/

Alignment-free methods implemented a Convolutional Neural Networks adopted from previous work with modifications. To identify the consensus features from different species, each sequence was first mapped to the feature space, ***F***, to create the feature vectors according to a finite alphabet, ***A***. After mapping, the similarity between feature vectors was calculated. We trained a simple CNN with a total of 8 hidden layers, including a one-dimensional convolution on top of the feature vectors. Consistent with the concept used in image processing, we also added zero padding around the feature vectors to get the same dimensions as input^40, 41^ so that no features would be shrinking. The maximum (Max) pooling layer was used in between to extract the sharpest features and reduce the complexity.^42-44^ We applied 20 sliding filters with different kernel sizes to optimize performance. To prevent co-adaptation of the hidden layers and make the model more robust, we apply regularization by employing dropout layers.^45, 46^ Twenty percent of the hidden units were dropped out. Last, we added a dense layer with one as parameter to produce a single output node in the output vector.^47, 48^ Sources and versions of open-source software libraries and neural-network library are listed below.

- NumPy v1.20.0 (https://numpy.org/)
- Pandas v1.2.4 (https://pandas.pydata.org/)
- TensorFlow v2 (https://www.tensorflow.org/install)
- Biopython v1.78 (https://biopython.org/wiki/Download)
- Matplotlib v3.4.2 (https://matplotlib.org/stable/users/installing.html)

Python scripts used in the study are found on GitHub: https://github.com/rhart604/optimized

## 3. Results

To generate ‘hypothetical’ RNAseq data with pooled reads from both human and mouse, we randomly selected a total of 9,500,000 paired-ended reads from two FASTQ files with different percentages of human content (0, 10, 50, 90 or 100%). To track the source of each read, a prefix of ‘human-’ or ‘mouse-’ was added to the read ID in the FASTQ records.

As an example of the potential for misrepresentation of alignment counts, we analyzed the 50% mixture of reads aligned with a mixed human and mouse reference genome using HISAT2 (**Table 1**). By comparing the read IDs pre-tagged with source genome vs. the species of the aligned reference chromosome, the total numbers of correctly paired alignments could be counted, along with a percent misalignment (the fraction of total aligned read pairs assigned to the wrong genome). While this fraction was small (0.15% for human reads and 0.40% for mouse reads), it was not zero. Furthermore, the numbers of pairs aligned exceeded the input reads from each species, by 4.43% for hg38 and 0.40% for mm10. More concerning was the observation that summary counts for several individual genes had a substantial misassignment due to human reads matching mouse genome (**Supplemental Table 1**). The top five genes on this list (**Table 2**) exhibited a surprisingly high proportion of misaligned reads, ranging from ∼8% to over 65%. Clearly, using a mixed reference genome allows for cross-alignment of reads to the wrong species, and for some genes this can produce a substantial source of error.

**Table 1.** HISAT2 alignment of a 50-50 mixture of human or mouse FASTQ reads with a mixed genome reference index. Counts are shown for first in pair properly aligned (flag 0×43) to each genome. The percent misaligned is the fraction of total read pairs from each species aligning with the incorrect genome.

| Source | hg38 | mm10 | Percent Misaligned | Percent False Positive |
| --- | --- | --- | --- | --- |
| human | 4,960,218 | 7,526 | 0.15% | 4.43% |
| mouse | 37,591 | 4,769,160 | 0.78% | 0.40% |
| Total read pairs aligned | 4,997,809 | 4,776,686 |  |  |

**Table 2.** Top 5 genes with the highest proportion of human reads matching mouse genome ortholog and at least 10 counts per genome. For complete list, see Supplemental Table 1.

| Mouse symbol | mm10 counts | Human symbol | hg38 counts | Percent mm10 |
| --- | --- | --- | --- | --- |
| Lars2 | 274 | LARS2 | 128 | 68.16% |
| Srsf1 | 106 | SRSF1 | 1009 | 9.51% |
| Rc3h2 | 26 | RC3H2 | 263 | 9.00% |
| Bcl11a | 15 | BCL11A | 167 | 8.24% |
| Clk4 | 17 | CLK4 | 212 | 7.42% |

Two general strategies were pursued—those that relied upon alignment with a reference genome index and those that did not. We will compare these along with minor variations to explore the optimal method for parsing mixed-genome samples.

### 3.1. Alignment-Dependent Methods

#### 3.1.1. HISAT2 with Separate Genome Index

The first method we tested aligned RNAseq reads with either the mouse reference genome index or the human reference genome index independently. Since results with a mixed genome index identified misalignments, we hypothesized that separately aligning with each genome would have fewer opportunities for mismatch based on homologous sequence.

Using **hisat2-build**, we created separate indices for the human genome and the mouse genome using toplevel fasta DNA sequence files from the Ensembl ftp site (GRCh38 [hg38] or GRCm38 [mm10], respectively). After building the indices, we used HISAT2 to align each of the 5 pairs of mixed fastq files with a range of species compositions using each index. The result was 5 SAM files for human and 5 for mouse. The number of non-matching and matching reads in each SAM file was recorded. As expected, some reads matched both species, and so were counted twice, inflating the total percent alignment above 100% (**Fig. 1A**, HISAT2_sep). The percent error was calculated as the sum of the differences between actual vs. observed proportion of reads for each species. The high number of non-matching reads contributed to an unexpectedly high percentage error, as shown in the “HISAT2_sep” bars in **Fig. 1B**. The accuracy, defined as the percentage of human reads aligned with human genome, was relatively insensitive to the species proportions (**Fig. 1C**). Separating the alignment produced greater total alignments with greater error and a similar accuracy, compared with mixed-genome alignment.

**Figure 1.**
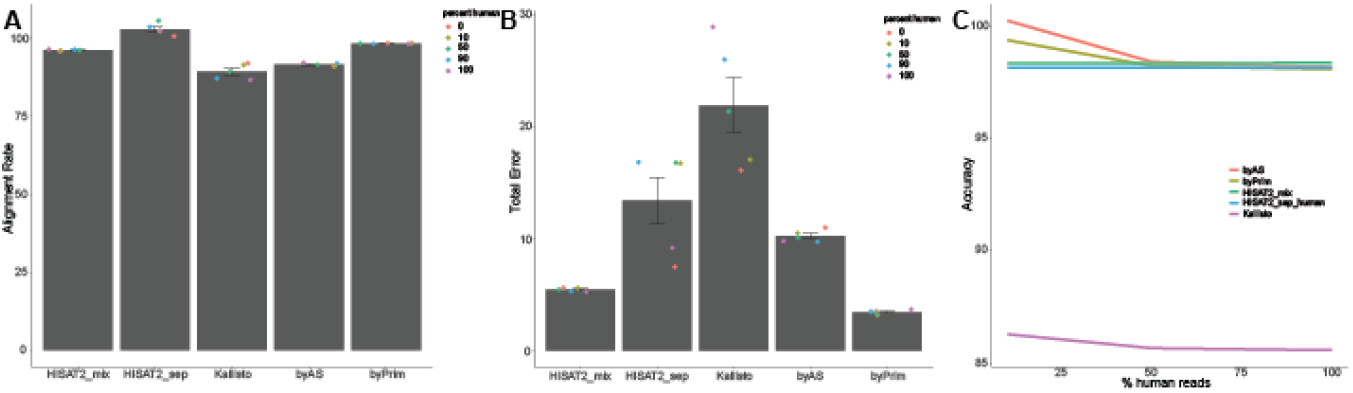
Comparison of alignment methods. A. Percentage of reads aligned using each method. Dots indicate the proportion of the two genomes. B. The error rate is calculated as the difference between fraction of reads aligned and the expected fraction, summed for each genome. C. Accuracy was assessed for each method and genome proportion by comparing the count of human read pairs correctly aligned to the human genome to the number of human read pairs in the input file.

#### 3.1.2. Kallisto with Combined Index

The high error using HISAT2 on separate genomes led us to consider using Kallisto, which uses pseudoalignment to efficiently quantify abundances of transcripts.^49^ Unlike HISAT2, which indexes the entire genome, Kallisto builds an index considering only subsequences, or k-mers, that distinguish isoforms. We built an index of the pooled mouse and human cDNA sequence, intending to focus on only the k-mers that distinguish the two different species (as well as different isoforms or genes within a species).

First, cDNA FASTA files from the Ensembl archive were downloaded from both GRCh38 and GRCm38, with each transcript identified by a species-specific Ensembl transcript ID. The files were combined and run through **kallisto index** to build an index. Then **kallisto quant** was used to align each of the 5 pairs of mixed files to the combined index. The number of mouse sequences that matched to human species and vice versa in the resulting directories were counted using the R package Rsubread (**Fig. 1**). Kallisto produced slightly fewer alignments than HISAT2 (third bar, **Fig. 1A**). While Kallisto was expected to be more accurate than the mixed-genome HISAT2 method, the average percent error was 11.2%, compared with only 1.7% for HISAT2-mixed (**Fig. 1B**). The accuracy, however, was substantially lower than other alignment methods (**Fig. 1C**), causing us to abandon this approach.

#### 3.1.3. Separating FASTQ Reads by Alignment Score

To improve on interpretation of alignments, the “by alignment score” (byAS) method was developed, based on the hypothesis that among multiple alignments from each sequencing record to the mixed genome index, the one with the greatest alignment score is the best match. These scores are used to split the original sequencing files into two sets of species-specific RNAseq reads for re-alignment with only the appropriate genome.

We prepared a Python script (byAS.py) that separates RNAseq FASTQ files into human and mouse subsets based on their alignment scores in SAM files produced from HISAT2 with a mixed reference index. The alignment score, a component of the SAM file format, is calculated by each algorithm using its own method, so it can only be compared among results from a particular program.

The byAS script scans a SAM file and stores alignment scores from each species in a separate dictionary, in which the key is the unique record ID. Reads that match both genomes with equal scores (∼0.51% of reads) are biased to choose the first-appearing read in the bam file. The two species-specific dictionaries are then used to iterate through the source, paired FASTQ files, exporting records into two sets of species-specific FASTQ files. The two sets of paired FASTQ files are re-aligned with the appropriate species of genome index using HISAT2. Selecting only high-scoring read pairs slightly reduced the final alignment rate (**Fig. 1A**, byAS). However, the overall error rate (**Fig. 1B**) is substantially reduced compared with a direct alignment with individual genomes (HISAT2_sep), and the accuracy was equal to or better than other methods (**Fig. 1C**). The strategy of classifying reads based on optimal match to a given genome clearly improved accurate alignment.

#### 3.1.4. Separating FASTQ Reads by Primary Alignment Flag

During evaluation of the byAS method, we realized that the alignment score calculated by HISAT2 exhibits a narrow range of values so there is little resolution between the best and worst alignments. Furthermore, the number of reads with identical alignment scores (∼0.51%), while small, is another source of inaccuracy. Another element of the SAM file format is a flag to denote the “primary” alignment, with lesser quality alignments considered as secondary (marked with a “true” SAM flag at 0×100, or decimal 256, which denotes “not primary alignment”). Since this flag, when set as “false,” ought to be found on the read with the best alignment score, it appeared to provide a simpler strategy for classifying alignments by species. We designed a second Python script (byPrim.py) to split FASTQ files based on the primary alignment flag in a mixed-genome SAM file.

This script reads the SAM file input and stores all read IDs as keys and primary flags as values in two species-specific dictionaries. Then, using these dictionaries, byPrim parses through the paired FASTQ files and outputs two sets of species-specific files.

Results indicated a slightly greater alignment rate (**Fig. 1A**, byPrim), reduced error rate (**Fig. 1B**), and a similar accuracy (**Fig. 1C**) compared with other HISAT2-based methods. However, the script is slightly simpler, with less ambiguity (for example, there are no reads excluded for matching alignment scores). Based on these evaluations, byPrim was judged to be optimal among the alignment methods.

### 3.2. Alignment-Independent Methods

The aim of an alignment-independent method is to build a classifier to distinguish sequence reads from different species without aligning to individual reference genomes. This requires us capture the hidden information directly from the nucleotide sequence of both mouse and human. We first sought to apply a classical probabilistic based approach, Hidden Markov models (HMMs), to discover the underlying variance between human and mouse sequence. HMMs are used in sequence data analysis with many bioinformatics applications^50, 51^ including identification of genes, motifs finding, metagenomic taxonomic classification.^52-54^ However, third order HMMs did not separate sequence fragments from mouse and human (**Fig. 2A**). Even with higher order Markov Models (8th or higher) which successfully performed metagenomic sequence classification,^55^ the separation of human and mouse reads is not ideal. The receiver operating characteristic curve^56^ indicates that this binary classifier system only slightly improves with higher order models (**Fig. 2B**). Moreover, we noticed that HMMs require substantial amount of memory and compute time. For a sequence of length ***l***, the memory to find the best path through the model with ***s*** states and ***e*** edges proportional to ***sl*** and the time proportional to ***el***.

**Figure 2.**
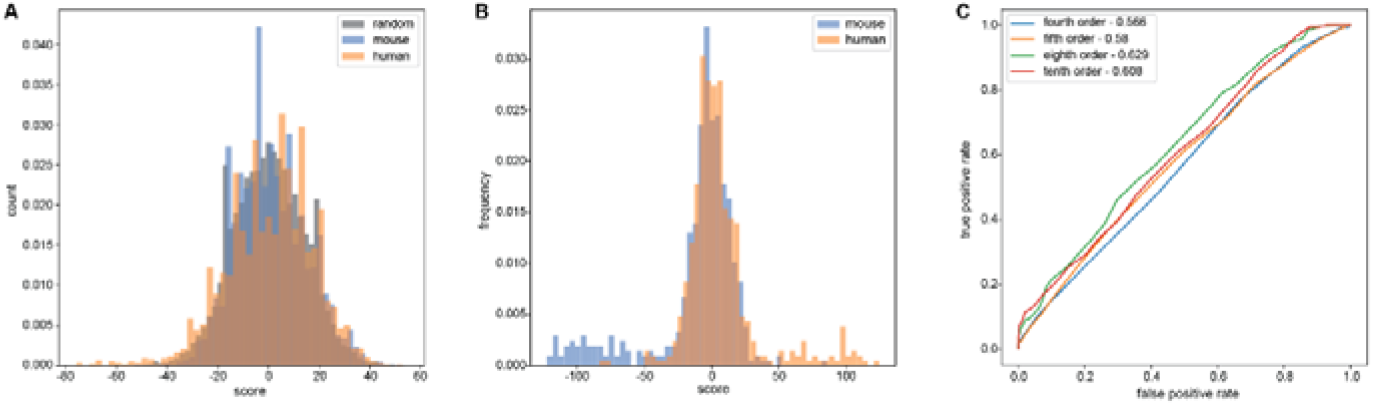
Performance of Hidden Markov Models. A. Separation of human and mouse reads compared with randomly generated sequence with same length using third order Markov Model. B. Separation of human and mouse reads with 10^th^ order of Markov Models. C. ROC plot showing the false positive rate and true positive rate of different orders of Markov models.

To find a more accurate and efficient way to separate human and mouse reads, we next utilized Convolutional Neural Networks (CNN),^47^ which was originally developed for computer visualization and image processing.^57, 58^ CNN performs effectively in many natural language processing (NLP) problems including sentence modeling, text classification and query processing.^42, 59-62^ We applied CNN here to classify RNA sequencing reads of different species using the identified features directly from the fragments without pre-aligning to reference genome.

We consider this process as assigning labels of either human or mouse to strings of characters. The key step is choosing the sets of features which will then input into the classification algorithms. Features like sequence length are highly dependent on the size selection step during library preparation so input strings must be truncated to a constant length (**Supplemental Fig. 1A**).^63^ Other sequence features, for example GC content, may have specificity for human genome.^64^ However, the difference in GC content between these two species (∼41% in human and ∼42% in mouse for non-sex chromosomes; **Supplemental Fig. 1B**) is insufficient to aid the separation.^65^

Our approach was to utilize raw reads from FASTQ files. Reads in the FASTQ file can be represented as a linear succession of L characters or nucleotides. Each nucleotide was coded using a finite alphabet, ***A***, containing five nucleotides, A, T, C, G, or N, pointing to integers 1,2,3,4, or 5, respectively, where ‘N’ is denoted as an ambiguous base due to low quality during sequencing.^66^ The percentage of N bases per read was calculated for each species to ensure there was no substantial difference (**Supplemental Fig. 1C**). All the generic strings can be obtained by concatenating characters from ***A*** to create the sample space, ***S***. Specifically, each sequencing read, *r*, can be mapped to feature space, ***F***, from sample space, ***S***, by a function : ℱ using the alphabetic index. We then represent each string *r* of length *l* as a multidimensional feature vector, *x*, in the 5^*1*^ dimensional feature space by***x*** = : ℱ (***r***) according to the alphabet table.

The basic architecture of the convolutional neural network is adopted from the original work of Collobert et al.^42^ in NLP with some modifications (**Supplemental Fig. 2G**). To avoid feature shrinking, each vector of length *l* was first padded with zeros to equalize dimensions before feeding into the network.^40, 41^ For each convolutional layer, we use a convolution operation with fixed filter width, *k*, to produce a new feature by given function *φ*. Here, *φ* is a non-linear function that may involve a bias term. The filter is applied to the entire feature vector with all possibilities, creating a new feature map. We then apply a max pooling operation over the feature map and take the maximum value as output from this layer.^42^ The penultimate layer takes all features obtained from the previous layers and outputs a probability distribution over either label. The last dense layer then outputs a binary classification indicating the species. Note that we also employed a dropout on the penultimate layer to avoid the co-adaptation of units.^45^ We introduce a random variable from a Bernoulli distribution, with probability *p* being 1, which determines the proportion of dropping units.

To test the performance of our model, we used the mixed human and mouse reads with different ratios. Reads are drawn from sequencing output without any selection or tuning. Reads were further split randomly into a 50% training set and a 50% testing set. We trained the same network with five training sets and tested the performance of each trained network using all five testing sets. The final performance was evaluated by accuracy after six epochs. Accuracies of model trained by either 0% or 100% human showed a strict linear relation with percentage of human reads in the testing sets (**Fig. 3A**). The 10% and 50% human training sets behaved with similar accuracies across all testing sets (**Fig. 3A**). With 50% or less human reads in the testing set, models that are trained by training sets with 10% and 50% human reads outperformed models trained by other training sets. With 90% or more human reads in the testing sets, models trained by 90% and 100% human reads performed similarly. The total accuracy positively correlated with the ratio in testing sets. To further determine the conceptual training strategy with different ratios of human reads in the testing sets, we compared accuracies with multiple possible testing datasets. The performance of models tested with 0, 10% and 50% human reads decrease along with decreased human reads in training sets (**Fig. 3B**). However, the accuracies dropped at 50% or more human reads in the training sets. This intuitively makes sense since more human reads in the training data would have emphasized the features of human rather than mouse which leads to the worse performance with lower human reads in the testing. The accuracy from testing sets with a higher percentage of human reads, on the other hand, jumped significantly after 10% then increased along with increased human reads in the training sets. Thus, we propose that to optimize performance of the model, the training sets should contain equal number of reads from both species.

**Figure 3.**
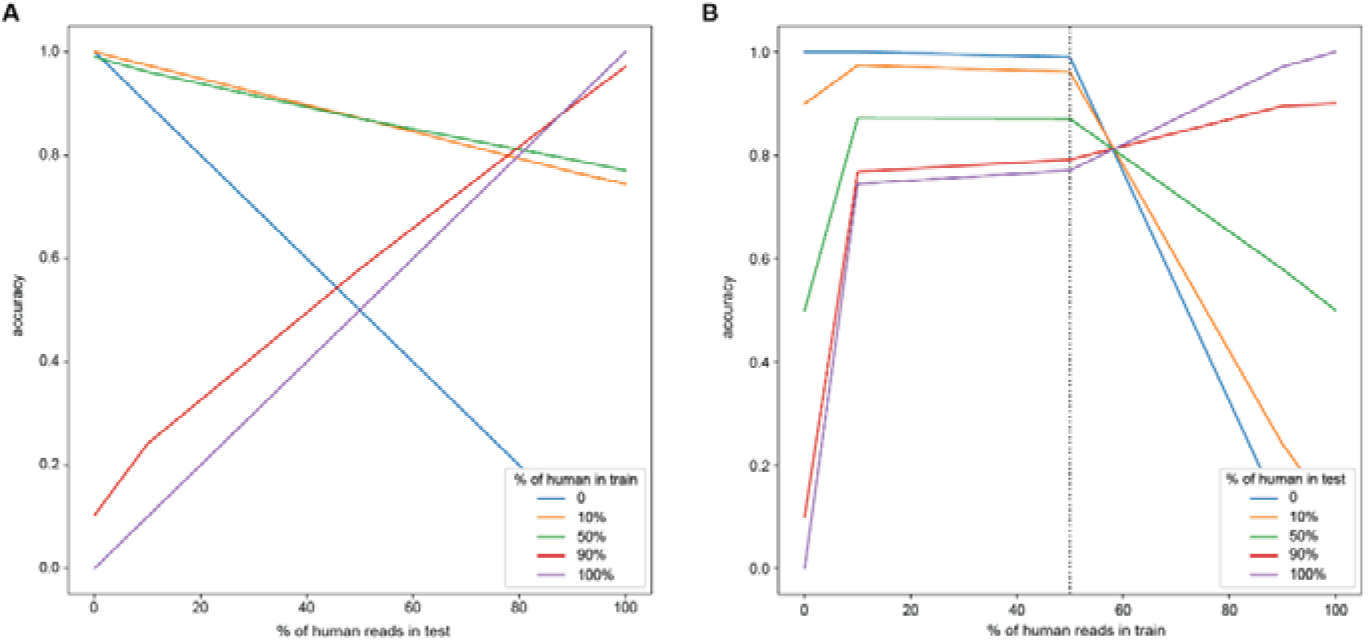
Performance of Convolutional Neural Networks. A. Accuracy of classification with different ratios of human reads in the training datasets. B. Accuracy of classification with different ratios of human reads in the testing datasets. Accuracy was calculated as the percentage of correctly classified reads in the input data.

To better classify human and mouse reads, we tuned several hyperparameters including kernel size, filters numbers, pooling methods and optimization algorithm to produce the highest accuracy (**Supplemental Fig. 2**). First, we noticed that the number of epochs with the highest accuracy turned out to be five, which is smaller than the usual number required in image processing or text classification. The accuracy on training sets increased dramatically after five epochs and continued increase to near 100% while the accuracy on testing sets dropped after five iterations (**Supplemental Fig. 2A**). This indicated a quick overfitting of the model^67, 68^ and signaled that the number of epochs should be set strictly within six.^69-71^ However, variations in the kernel size and number of filters, which are two critical parameters that alter network efficiency,^72, 73^ did not change the classification accuracy dramatically (**Supplemental Fig. 2B-C**). Kernel size only affected the overall performance of the model by around 1%. With a kernel size larger than four, the model stably produces accuracies over 87% (**Supplemental Fig. 2B-C**). Similarly, filter number only slightly changed the accuracy (**Supplemental Fig. 2D**). Pooling layer, on the other hand, was crucial after the convolution layer. Not only did it downsize the information to reduce computation time, but it also increased the classification accuracy.^74^ Network without pooling usually had a substantial decrease of performance which was usually caused by the propagation of local features to the neighbors.^75^ So, including the pooling operation can shrink the feature map while still preserving key information required for classification.^43, 76, 77^ We investigated two popular pooling methods, average pooling and max pooling, each with its own strengths and weaknesses.^43, 78^ The evaluation of the model accuracy with different pooling methods showed that max pooling outperformed average pooling despite the percentage of human reads (**Supplemental Fig. 2E**). Finally, we showed that instead of classical stochastic gradient descent procedure, the Adam optimization algorithm,^79-81^ performed the best (**Supplemental Fig. 2F**).

## 4. Discussion

Our results demonstrate that the optimal strategy for obtaining accurate classification of sequencing reads from mixed-species samples requires three steps: first, alignment to a mixed reference genome, then, the selection of the optimal alignment for each read to partition reads into species-specific FASTQ files, followed by re-alignment to the appropriate genome. Aligning mixed samples using a pooled reference index resulted in errors, some of which underestimated counts unequally across genes. Surprisingly, an algorithm designed to focus on differences within a reference sequence (Kallisto) was less effective at distinguishing reads by species. An algorithm using indexed genomes (HISAT2) performed better, with better overall accuracy and reduced error, particularly when coupled with a method to separate input FASTQ files using alignment information. With pooled mouse and human RNAseq reads with various proportions, both byAS and byPrim methods offered over 95% accuracies.

The overall performance of alignment-based method, however, is highly dependent on the quality of the sequencing data, and could be less effective under conditions where poor sequencing quality leads to lower alignment rate. Therefore, we also evaluated alignment-independent methods, HMM and CNN, by which reads can be separated based on features within the sequences without pre-aligning to reference genomes. HMM did not adequately separate reads by species. However, CNN could be applied for sequencing datasets from organisms whose genomes are not well annotated. Importantly, CNN provides better and faster classification of RNAseq reads of two species compared with HMM (**Fig. 2**). We suspect the suboptimal performance of such probabilistic models is due to the high similarity between the linear sequence from human and mouse genome.

In summary, while the optimal CNN non-alignment strategy successfully partitioned reads by species, a more traditional approach of mixed-genome alignment followed by separation of reads by optimal alignment (byPrim) proved to be the most successful with the lowest error rates.

## 5. Acknowledgments

*Supported by grants from NIH (U10AA008401, R01ES026057) and startup funds from the University of Pittsburgh to KH. The authors acknowledge the Office of Advanced Research Computing (OARC) at Rutgers, The State University of New Jersey for providing access to the Amarel cluster and associated research computing resources*.

## Supplemental Figures

**Supplemental Figure 1.**
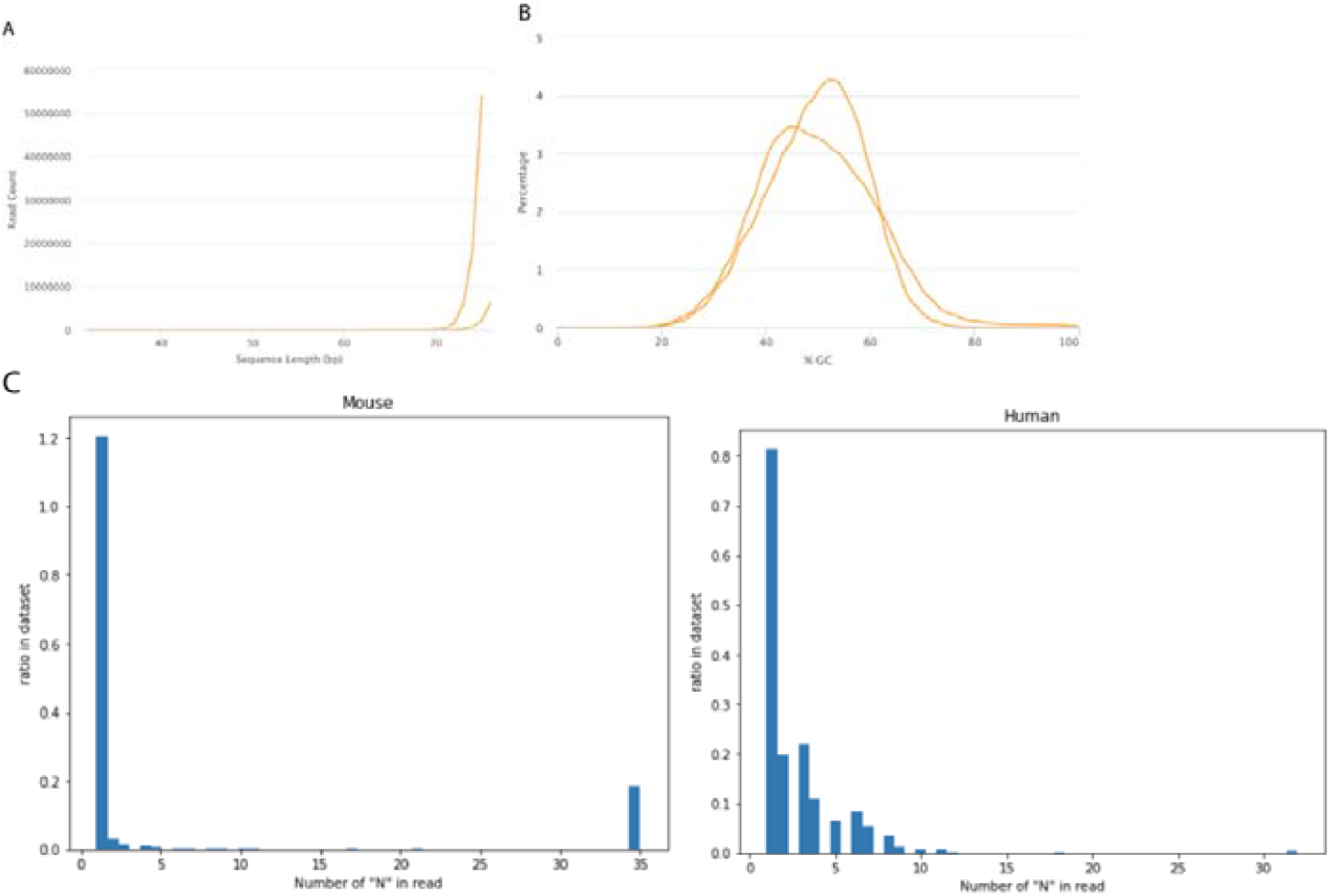
Characteristic of sequence reads. A. Length distribution of human and mouse reads. B. GC content of human and mouse reads. C. Ratio of ambiguous base ‘N’ in human and mouse sequence.

**Supplemental Figure 2.**
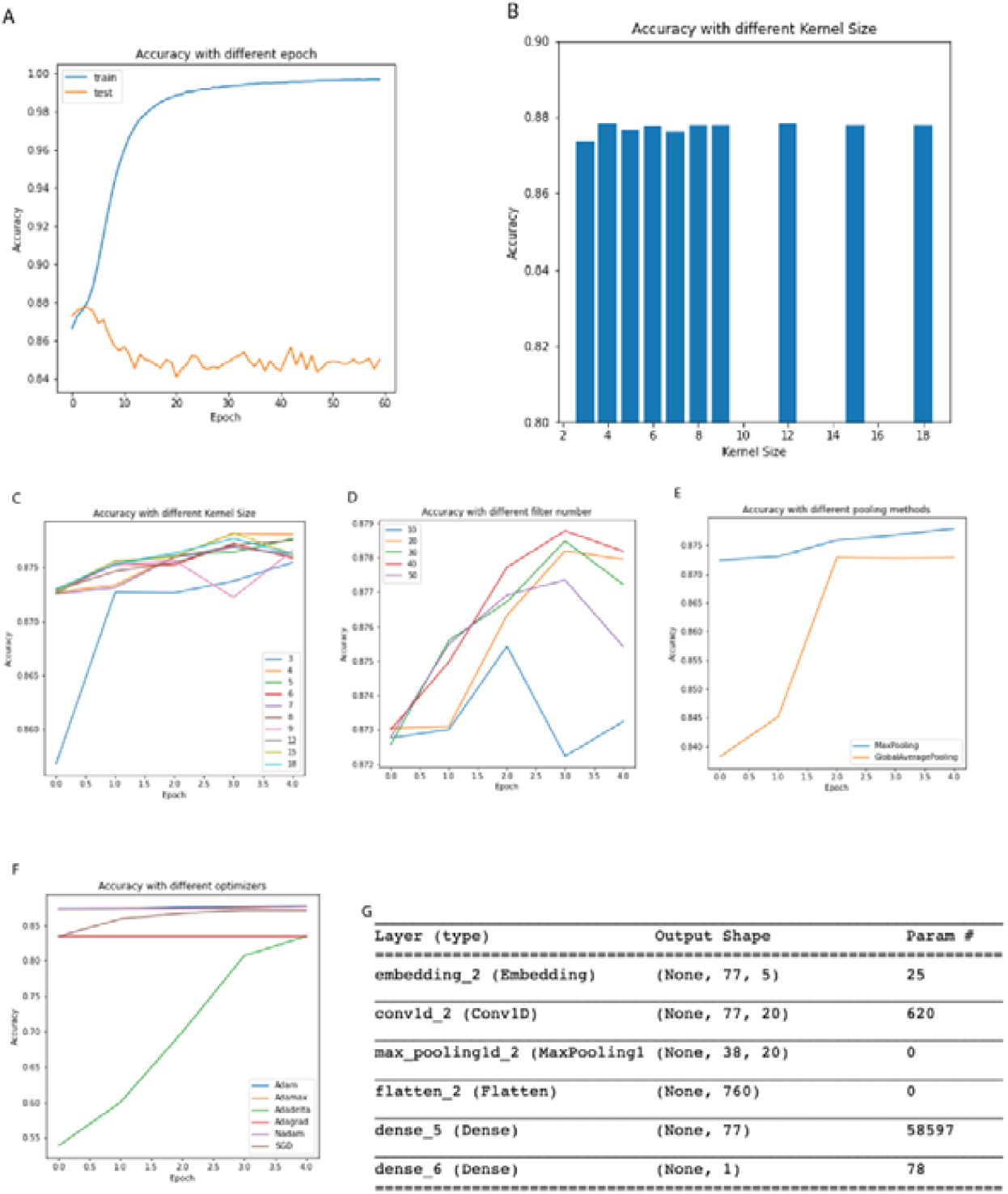
Optimization of hyperparameters. A. Accuracy of classification in training and testing sets with increase of epochs. B. Comparison of accuracy of classification with different kernel size using six epochs. C. Accuracy of classification with different kernel size during first five epochs. D. Comparison of accuracy of classification with different filter numbers using six epochs. E. Comparison of accuracy of classification with different pooling methods. F. Comparison of accuracy of classification with different optimizers. Accuracy was calculated as the percentage of correctly classified reads in the input data. G. Structure of Neural Networks.

## Notes

https://github.com/rhart604/optimized

### Competing Interest Statement

The authors have declared no competing interest.

## References

1. Morse III H. C. 2007, “Building a better mouse: One hundred years of genetics and biology,” in The mouse in biomedical research, ed.^eds. Editior (Elsevier, pp. Elsevier.

2. Rosenthal N. and Brown S. 2007, “The mouse ascending: perspectives for human-disease models,” Nature cell biology 9, 993–999.

3. Waterston R. H. and Pachter L. 2002, “Initial sequencing and comparative analysis of the mouse genome,” Nature 420, 520–562.

4. Bedell M. A., Jenkins N. A. and Copeland N. G. 1997, “Mouse models of human disease. Part I: techniques and resources for genetic analysis in mice,” Genes & development 11, 1–10.

5. Hoyt R., Hawkins J., St Clair M. and Kennett M. 2007, “The mouse in biomedical research,” American College of Laboratory Animal Medicine 3.

6. Kerbel R. S. 1998, “What is the optimal rodent model for anti-tumor drug testing?,” Cancer and Metastasis Reviews 17, 301–304.

7. Elsea S. H. and Lucas R. E. 2002, “The mousetrap: what we can learn when the mouse model does not mimic the human disease,” ILAR journal 43, 66–79.

8. Arranz A. M., Espuny Camacho I. M., Fiers M., Snellinx A., Ando K., Munck S., Corthout N., Radaelli E., Leroy K. and Brion J. P. 2017, “Hallmarks of Alzheimer’s disease in stem cell-derived human neurons transplanted into mouse brain,” in GLIA, ed.^eds. Editior (WILEY, pp. WILEY.

9. Crews L. and Masliah E. 2010, “Molecular mechanisms of neurodegeneration in Alzheimer’s disease,” Human molecular genetics 19, R12–R20.

10. Lin S., Lin Y., Nery J. R., Urich M. A., Breschi A., Davis C. A., Dobin A., Zaleski C., Beer M. A. and Chapman W. C. 2014, “Comparison of the transcriptional landscapes between human and mouse tissues,” Proceedings of the National Academy of Sciences 111, 17224–17229.

11. Thapa K. S., Chen A. B., Lai D., Xuei X., Wetherill L., Tischfield J. A., Liu Y. and Edenberg H. J. 2020, “Identification of Functional Genetic Variants Associated With Alcohol Dependence and Related Phenotypes Using a High-Throughput Assay,” Alcohol Clin Exp Res 44, 2494–2518.

12. Xiao X., Chang H. and Li M. 2017, “Molecular mechanisms underlying noncoding risk variations in psychiatric genetic studies,” Mol Psychiatry 22, 497–511.

13. Bocher O. and Génin E. 2020, “Rare variant association testing in the non-coding genome,” Hum Genet 139, 1345–1362.

14. Low L. K. and Cheng H.-J. 2006, “Axon pruning: an essential step underlying the developmental plasticity of neuronal connections,” Philosophical Transactions of the Royal Society B: Biological Sciences 361, 1531–1544.

15. Windrem M. S., Schanz S. J., Morrow C., Munir J., Chandler-Militello D., Wang S. and Goldman S. A. 2014, “A competitive advantage by neonatally engrafted human glial progenitors yields mice whose brains are chimeric for human glia,” Journal of Neuroscience 34, 16153–16161.

16. Real R., Peter M., Trabalza A., Khan S., Smith M. A., Dopp J., Barnes S. J., Momoh A., Strano A. and Volpi E. 2018, “In vivo modeling of human neuron dynamics and Down syndrome,” Science 362.

17. Espuny-Camacho I., Arranz A. M., Fiers M., Snellinx A., Ando K., Munck S., Bonnefont J., Lambot L., Corthout N. and Omodho L. 2017, “Hallmarks of Alzheimer’s disease in stem-cell-derived human neurons transplanted into mouse brain,” Neuron 93, 1066-1081. e8.

18. Mascetti V. L. and Pedersen R. A. 2016, “Human-mouse chimerism validates human stem cell pluripotency,” Cell stem cell 18, 67–72.

19. Xu R., Li X., Boreland A. J., Posyton A., Kwan K., Hart R. P. and Jiang P. 2020, “Human iPSC-derived mature microglia retain their identity and functionally integrate in the chimeric mouse brain,” Nature communications 11, 1–16.

20. Thompson L. H. and Björklund A. 2015, “Reconstruction of brain circuitry by neural transplants generated from pluripotent stem cells,” Neurobiology of disease 79, 28–40.

21. Shi Y., Kirwan P., Smith J., Robinson H. P. and Livesey F. J. 2012, “Human cerebral cortex development from pluripotent stem cells to functional excitatory synapses,” Nature neuroscience 15, 477–486.

22. Halikere A., Popova D., Scarnati M. S., Hamod A., Swerdel M. R., Moore J. C., Tischfield J. A., Hart R. P. and Pang Z. P. 2019, “Addiction associated N40D mu-opioid receptor variant modulates synaptic function in human neurons,” in Mol Psychiatry, ed.^eds. Editior.

23. Oni E. N., Halikere A., Li G., Toro-Ramos A. J., Swerdel M. R., Verpeut J. L., Moore J. C., Bello N. T., Bierut L. J., Goate A., Tischfield J. A., Pang Z. P. and Hart R. P. 2016, “Increased nicotine response in iPSC-derived human neurons carrying the CHRNA5 N398 allele,” Sci Rep 6, 34341.

24. Vierbuchen T., Ostermeier A., Pang Z. P., Kokubu Y., Sudhof T. C. and Wernig M. 2010, “Direct conversion of fibroblasts to functional neurons by defined factors,” Nature 463, 1035–41.

25. Pang Z. P., Yang N., Vierbuchen T., Ostermeier A., Fuentes D. R., Yang T. Q., Citri A., Sebastiano V., Marro S., Sudhof T. C. and Wernig M. 2011, “Induction of human neuronal cells by defined transcription factors,” Nature.

26. Fridman W. H., Pages F., Sautes-Fridman C. and Galon J. 2012, “The immune contexture in human tumours: impact on clinical outcome,” Nature Reviews Cancer 12, 298–306.

27. Rahier J., Goebbels R. and Henquin J.-C. 1983, “Cellular composition of the human diabetic pancreas,” Diabetologia 24, 366–371.

28. Wang J., Huang M., Torre E., Dueck H., Shaffer S., Murray J., Raj A., Li M. and Zhang N. R. 2018, “Gene expression distribution deconvolution in single-cell RNA sequencing,” Proceedings of the National Academy of Sciences 115, E6437–E6446.

29. Mohammadi S., Zuckerman N., Goldsmith A. and Grama A. 2016, “A critical survey of deconvolution methods for separating cell types in complex tissues,” Proceedings of the IEEE 105, 340–366.

30. Newman A. M., Liu C. L., Green M. R., Gentles A. J., Feng W., Xu Y., Hoang C. D., Diehn M. and Alizadeh A. A. 2015, “Robust enumeration of cell subsets from tissue expression profiles,” Nature methods 12, 453–457.

31. Baron M., Veres A., Wolock S. L., Faust A. L., Gaujoux R., Vetere A., Ryu J. H., Wagner B. K., Shen-Orr S. S. and Klein A. M. 2016, “A single-cell transcriptomic map of the human and mouse pancreas reveals inter-and intra-cell population structure,” Cell systems 3, 346-360. e4.

32. Wang X., Park J., Susztak K., Zhang N. R. and Li M. 2019, “Bulk tissue cell type deconvolution with multi-subject single-cell expression reference,” Nature communications 10, 1–9.

33. Ziegenhain C., Vieth B., Parekh S., Reinius B., Guillaumet-Adkins A., Smets M., Leonhardt H., Heyn H., Hellmann I. and Enard W. 2017, “Comparative analysis of single-cell RNA sequencing methods,” Molecular cell 65, 631-643. e4.

34. Jew B., Alvarez M., Rahmani E., Miao Z., Ko A., Garske K. M., Sul J. H., Pietiläinen K. H., Pajukanta P. and Halperin E. 2020, “Accurate estimation of cell composition in bulk expression through robust integration of single-cell information,” Nature communications 11, 1–11.

35. Wan X., Song H., Luo L., Li Z., Sheng G. and Jiang X. 2018, “Pattern recognition of partial discharge image based on one-dimensional convolutional neural network,” in 2018 Condition Monitoring and Diagnosis (CMD), ed.^eds. Editior (IEEE, pp. IEEE.

36. Kalchbrenner N., Espeholt L., Simonyan K., Oord A. v. d., Graves A. and Kavukcuoglu K. 2016, “Neural machine translation in linear time,” arXiv preprint 1610.10099.

37. Scarnati M. S., Boreland A. J., Joel M., Hart R. P. and Pang Z. P. 2020, “Differential sensitivity of human neurons carrying μ opioid receptor (MOR) N40D variants in response to ethanol,” Alcohol.

38. Shao Z., Noh H., Bin Kim W., Ni P., Nguyen C., Cote S. E., Noyes E., Zhao J., Parsons T., Park J. M., Zheng K., Park J. J., Coyle J. T., Weinberger D. R., Straub R. E., Berman K. F., Apud J., Ongur D., Cohen B. M., McPhie D. L., Rapoport J. L., Perlis R. H., Lanz T. A., Xi H. S., Yin C., Huang W., Hirayama T., Fukuda E., Yagi T., Ghosh S., Eggan K. C., Kim H. Y., Eisenberg L. M., Moghadam A. A., Stanton P. K., Cho J. H. and Chung S. 2019, “Dysregulated protocadherin-pathway activity as an intrinsic defect in induced pluripotent stem cell-derived cortical interneurons from subjects with schizophrenia,” Nat Neurosci 22, 229–242.

39. Clark S. C., Chereji R. V., Lee P. R., Fields R. D. and Clark D. J. 2020, “Differential nucleosome spacing in neurons and glia,” Neurosci Lett 714, 134559.

40. Albawi S., Mohammed T. A. and Al-Zawi S. 2017, “Understanding of a convolutional neural network,” in 2017 International Conference on Engineering and Technology (ICET), ed.^eds. Editior (Ieee, pp. Ieee.

41. Oquab M., Bottou L., Laptev I. and Sivic J. 2014, “Learning and transferring mid-level image representations using convolutional neural networks,” in Proceedings of the IEEE conference on computer vision and pattern recognition, ed.^eds. Editior.

42. Collobert R., Weston J., Bottou L., Karlen M., Kavukcuoglu K. and Kuksa P. 2011, “Natural language processing (almost) from scratch,” Journal of machine learning research 12, 2493− 2537.

43. Tolias G., Sicre R. and Jégou H. 2015, “Particular object retrieval with integral max-pooling of CNN activations,” arXiv preprint 1511.05879.

44. Nagi J., Ducatelle F., Di Caro G. A., Cireşan D., Meier U., Giusti A., Nagi F., Schmidhuber J. and Gambardella L. M. 2011, “Max-pooling convolutional neural networks for vision-based hand gesture recognition,” in 2011 IEEE International Conference on Signal and Image Processing Applications (ICSIPA), ed.^eds. Editior (IEEE, pp. IEEE.

45. Hinton G. E., Srivastava N., Krizhevsky A., Sutskever I. and Salakhutdinov R. R. 2012, “Improving neural networks by preventing co-adaptation of feature detectors,” arXiv preprint 1207.0580.

46. Poernomo A. and Kang D.-K. 2018, “Biased dropout and crossmap dropout: learning towards effective dropout regularization in convolutional neural network,” Neural networks 104, 60–67.

47. Husain S. S. and Bober M. 2019, “REMAP: Multi-layer entropy-guided pooling of dense CNN features for image retrieval,” IEEE Transactions on Image Processing 28, 5201–5213.

48. Zhang Y., Tian Y., Kong Y., Zhong B. and Fu Y. 2018, “Residual dense network for image super-resolution,” in Proceedings of the IEEE conference on computer vision and pattern recognition, ed.^eds. Editior.

49. Bray N. L., Pimentel H., Melsted P. and Pachter L. 2016, “Near-optimal probabilistic RNA-seq quantification,” Nat Biotechnol 34, 525–7.

50. Eddy S. R. 1998, “Profile hidden Markov models,” Bioinformatics (Oxford, England) 14, 755–763.

51. Munch K. and Krogh A. 2006, “Automatic generation of gene finders for eukaryotic species,” BMC bioinformatics 7, 1–12.

52. Wheeler T. J. and Eddy S. R. 2013, “nhmmer: DNA homology search with profile HMMs,” Bioinformatics 29, 2487–2489.

53. Finn R. D., Coggill P., Eberhardt R. Y., Eddy S. R., Mistry J., Mitchell A. L., Potter S. C., Punta M., Qureshi M. and Sangrador-Vegas A. 2016, “The Pfam protein families database: towards a more sustainable future,” Nucleic acids research 44, D279–D285.

54. Finn R. D., Clements J. and Eddy S. R. 2011, “HMMER web server: interactive sequence similarity searching,” Nucleic acids research 39, W29–W37.

55. Burks D. J. and Azad R. K. 2020, “Higher-order Markov models for metagenomic sequence classification,” Bioinformatics 36, 4130–4136.

56. Zou K. H., O’Malley A. J. and Mauri L. 2007, “Receiver-operating characteristic analysis for evaluating diagnostic tests and predictive models,” Circulation 115, 654–657.

57. Valueva M. V., Nagornov N., Lyakhov P. A., Valuev G. V. and Chervyakov N. 2020, “Application of the residue number system to reduce hardware costs of the convolutional neural network implementation,” Mathematics and Computers in Simulation 177, 232–243.

58. Behnke S. 2003, Hierarchical neural networks for image interpretation. (Springer).

59. Zhang Y. and Wallace B. 2015, “A sensitivity analysis of (and practitioners’ guide to) convolutional neural networks for sentence classification,” arXiv preprint 1510.03820.

60. Yih W.-t., Toutanova K., Platt J. C. and Meek C. 2011, “Learning discriminative projections for text similarity measures,” in Proceedings of the fifteenth conference on computational natural language learning, ed.^eds. Editior.

61. Shen Y., He X., Gao J., Deng L. and Mesnil G. 2014, “Learning semantic representations using convolutional neural networks for web search,” in Proceedings of the 23rd international conference on world wide web, ed.^eds. Editior.

62. Kalchbrenner N., Grefenstette E. and Blunsom P. 2014, “A convolutional neural network for modelling sentences,” arXiv preprint 1404.2188.

63. Kathiresan N., Temanni R., Almabrazi H., Syed N., Jithesh P. V. and Al-Ali R. 2017, “Accelerating next generation sequencing data analysis with system level optimizations,” Scientific reports 7, 1–11.

64. Piovesan A., Pelleri M. C., Antonaros F., Strippoli P., Caracausi M. and Vitale L. 2019, “On the length, weight and GC content of the human genome,” BMC research notes 12, 1–7.

65. Guénet J. 2005, “9 Inducing Alterations in the Mammalian Genome for Investigating the Functions of Genes,” Mammalian Genomics, 221.

66. Ewing B., Hillier L., Wendl M. C. and Green P. 1998, “Base-calling of automated sequencer traces usingPhred. I. Accuracy assessment,” Genome research 8, 175–185.

67. Gavrilov A. D., Jordache A., Vasdani M. and Deng J. 2018, “Preventing model overfitting and underfitting in convolutional neural networks,” International Journal of Software Science and Computational Intelligence (IJSSCI) 10, 19–28.

68. Özgenel Ç. F. and Sorguç A. G. 2018, “Performance comparison of pretrained convolutional neural networks on crack detection in buildings,” in ISARC. Proceedings of the International Symposium on Automation and Robotics in Construction, ed.^eds. Editior (IAARC Publications, pp. IAARC Publications.

69. Arif R. B., Siddique M. A. B., Khan M. M. R. and Oishe M. R. 2018, “Study and Observation of the Variations of Accuracies for Handwritten Digits Recognition with Various Hidden Layers and Epochs using Convolutional Neural Network,” in 2018 4th International Conference on Electrical Engineering and Information & Communication Technology (iCEEiCT), ed.^eds. Editior (IEEE, pp. IEEE.

70. Phan H., Andreotti F., Cooray N., Chén O. Y. and De Vos M. 2018, “Joint classification and prediction CNN framework for automatic sleep stage classification,” IEEE Transactions on Biomedical Engineering 66, 1285–1296.

71. Zhu X. and Bain M. 2017, “B-CNN: branch convolutional neural network for hierarchical classification,” arXiv preprint 1709.09890.

72. Agrawal A. and Mittal N. 2020, “Using CNN for facial expression recognition: a study of the effects of kernel size and number of filters on accuracy,” The Visual Computer 36, 405–412.

73. Huang Z., Dong M., Mao Q. and Zhan Y. 2014, “Speech emotion recognition using CNN,” in Proceedings of the 22nd ACM international conference on Multimedia, ed.^eds. Editior.

74. Akhtar N. and Ragavendran U. 2020, “Interpretation of intelligence in CNN-pooling processes: a methodological survey,” Neural Computing and Applications 32, 879–898.

75. Springenberg J. T., Dosovitskiy A., Brox T. and Riedmiller M. 2014, “Striving for simplicity: The all convolutional net,” arXiv preprint 1412.6806.

76. Gong Y., Wang L., Guo R. and Lazebnik S. 2014, “Multi-scale orderless pooling of deep convolutional activation features,” in European conference on computer vision, ed.^eds. Editior (Springer, pp. Springer.

77. Chen J., Hua Z., Wang J. and Cheng S. 2017, “A convolutional neural network with dynamic correlation pooling,” in 2017 13th International Conference on Computational Intelligence and Security (CIS), ed.^eds. Editior (IEEE, pp. IEEE.

78. Xu Z., Yang Y. and Hauptmann A. G. 2015, “A discriminative CNN video representation for event detection,” in Proceedings of the IEEE conference on computer vision and pattern recognition, ed.^eds. Editior.

79. Koushik J. and Hayashi H. 2016, “Improving stochastic gradient descent with feedback.”

80. Kingma D. P. and Ba J. 2014, “Adam: A method for stochastic optimization,” arXiv preprint 1412.6980.

81. Reddi S. J., Kale S. and Kumar S. 2019, “On the convergence of adam and beyond,” arXiv preprint

