## Supplemental Table 1 for "Optimized splitting of RNA sequencing data by species"

| mm10 symbol | mm10 counts | mm10 ensembl | hg38 ensembl | hgnc symbol | hg38 counts | Percent |
|---|---|---|---|---|---|---|
| Lars2 | 274 | ENSMUSG00000035202 | ENSG00000011376 | LARS2 | 128 | 68.16% |
| Srsf1 | 106 | ENSMUSG00000018379 | ENSG00000136450 | SRSF1 | 1009 | 9.51% |
| Rc3h2 | 26 | ENSMUSG00000075376 | ENSG00000056586 | RC3H2 | 263 | 9.00% |
| Bcl11a | 15 | ENSMUSG00000000861 | ENSG00000119866 | BCL11A | 167 | 8.24% |
| Clk4 | 17 | ENSMUSG00000020385 | ENSG00000113240 | CLK4 | 212 | 7.42% |
| Cnot2 | 35 | ENSMUSG00000020166 | ENSG00000111596 | CNOT2 | 467 | 6.97% |
| Fmr1 | 13 | ENSMUSG00000000838 | ENSG00000102081 | FMR1 | 200 | 6.10% |
| Nova1 | 30 | ENSMUSG00000021047 | ENSG00000139910 | NOVA1 | 468 | 6.02% |
| Srsf10 | 36 | ENSMUSG00000028676 | ENSG00000188529 | SRSF10 | 582 | 5.83% |
| Srsf3 | 72 | ENSMUSG00000071172 | ENSG00000112081 | SRSF3 | 1299 | 5.25% |
| Matr3 | 195 | ENSMUSG00000037236 | ENSG00000015479 | MATR3 | 3530 | 5.23% |
| Tra2b | 43 | ENSMUSG00000022858 | ENSG00000136527 | TRA2B | 870 | 4.71% |
| Nrxn3 | 15 | ENSMUSG00000066392 | ENSG00000021645 | NRXN3 | 310 | 4.62% |
| Prpf39 | 15 | ENSMUSG00000035597 | ENSG00000185246 | PRPF39 | 348 | 4.13% |
| Hnrnph1 | 96 | ENSMUSG00000007850 | ENSG00000169045 | HNRNPH1 | 2333 | 3.95% |
| Eif1 | 96 | ENSMUSG00000035530 | ENSG00000173812 | EIF1 | 2475 | 3.73% |
| Hnrnph3 | 42 | ENSMUSG00000020069 | ENSG00000096746 | HNRNPH3 | 1094 | 3.70% |
| Zbtb18 | 12 | ENSMUSG00000063659 | ENSG00000179456 | ZBTB18 | 347 | 3.34% |
| Tra2a | 16 | ENSMUSG00000029817 | ENSG00000164548 | TRA2A | 469 | 3.30% |
| Pum1 | 24 | ENSMUSG00000028580 | ENSG00000134644 | PUM1 | 705 | 3.29% |
| Srsf11 | 37 | ENSMUSG00000055436 | ENSG00000116754 | SRSF11 | 1117 | 3.21% |
| Hnrnpa3 | 42 | ENSMUSG00000059005 | ENSG00000170144 | HNRNPA3 | 1358 | 3.00% |
| Cul3 | 14 | ENSMUSG00000004364 | ENSG00000036257 | CUL3 | 460 | 2.95% |
| Elavl4 | 17 | ENSMUSG00000028546 | ENSG00000162374 | ELAVL4 | 590 | 2.80% |
| Dhx15 | 25 | ENSMUSG00000029169 | ENSG00000109606 | DHX15 | 882 | 2.76% |
| Gria2 | 29 | ENSMUSG00000033981 | ENSG00000120251 | GRIA2 | 1034 | 2.73% |
| Srsf6 | 38 | ENSMUSG00000016921 | ENSG00000124193 | SRSF6 | 1360 | 2.72% |
| Fhl1 | 51 | ENSMUSG00000023092 | ENSG00000022267 | FHL1 | 1918 | 2.59% |
| Aph1a | 20 | ENSMUSG00000015750 | ENSG00000117362 | APH1A | 753 | 2.59% |
| Hnrnpr | 28 | ENSMUSG00000066037 | ENSG00000125944 | HNRNPR | 1121 | 2.44% |
| Ptbp2 | 15 | ENSMUSG00000028134 | ENSG00000117569 | PTBP2 | 617 | 2.37% |
| Ddx5 | 180 | ENSMUSG00000020719 | ENSG00000108654 | DDX5 | 7566 | 2.32% |
| Pfn2 | 38 | ENSMUSG00000027805 | ENSG00000070087 | PFN2 | 1619 | 2.29% |
| Atxn2l | 16 | ENSMUSG00000032637 | ENSG00000168488 | ATXN2L | 684 | 2.29% |
| Ogt | 19 | ENSMUSG00000034160 | ENSG00000147162 | OGT | 841 | 2.21% |
| Srsf7 | 14 | ENSMUSG00000024097 | ENSG00000115875 | SRSF7 | 620 | 2.21% |
| Rbmx | 37 | ENSMUSG00000031134 | ENSG00000147274 | RBMX | 1649 | 2.19% |
| Arglu1 | 24 | ENSMUSG00000040459 | ENSG00000134884 | ARGLU1 | 1101 | 2.13% |
| Luc7l3 | 45 | ENSMUSG00000020863 | ENSG00000108848 | LUC7L3 | 2247 | 1.96% |
| Fubp1 | 16 | ENSMUSG00000028034 | ENSG00000162613 | FUBP1 | 829 | 1.89% |
| Mat2a | 34 | ENSMUSG00000053907 | ENSG00000168906 | MAT2A | 1783 | 1.87% |
| Hnrnpdl | 58 | ENSMUSG00000029328 | ENSG00000152795 | HNRNPDL | 3080 | 1.85% |
| Paxbp1 | 14 | ENSMUSG00000022974 | ENSG00000159086 | PAXBP1 | 791 | 1.74% |
| Phc2 | 14 | ENSMUSG00000028796 | ENSG00000134686 | PHC2 | 834 | 1.65% |
| Son | 38 | ENSMUSG00000022961 | ENSG00000159140 | SON | 2264 | 1.65% |
| Pnn | 22 | ENSMUSG00000020994 | ENSG00000100941 | PNN | 1362 | 1.59% |
| Eif5 | 22 | ENSMUSG00000021282 | ENSG00000100664 | EIF5 | 1428 | 1.52% |
| Hmgb1 | 30 | ENSMUSG00000066551 | ENSG00000189403 | HMGB1 | 1953 | 1.51% |
| Syncrip | 13 | ENSMUSG00000032423 | ENSG00000135316 | SYNCRIP | 861 | 1.49% |
| Hnrnpk | 75 | ENSMUSG00000021546 | ENSG00000165119 | HNRNPK | 5184 | 1.43% |
| Nkx2-1 | 21 | ENSMUSG00000001496 | ENSG00000136352 | NKX2-1 | 1467 | 1.41% |
| Ppp1cc | 11 | ENSMUSG00000004455 | ENSG00000186298 | PPP1CC | 789 | 1.38% |
| Celf1 | 11 | ENSMUSG00000005506 | ENSG00000149187 | CELF1 | 806 | 1.35% |

| mm10 symbol | mm10 counts | mm10 ensembl | hg38 ensembl | hgnc symbol | hg38 counts | Percent |
|---|---|---|---|---|---|---|
| Ddx3x | 23 | ENSMUSG00000000787 | ENSG00000215301 | DDX3X | 1713 | 1.32% |
| Hnrnpu | 32 | ENSMUSG00000039630 | ENSG00000153187 | HNRNPU | 2393 | 1.32% |
| Ewsr1 | 19 | ENSMUSG00000009079 | ENSG00000182944 | EWSR1 | 1423 | 1.32% |
| Wsb1 | 22 | ENSMUSG00000017677 | ENSG00000109046 | WSB1 | 1658 | 1.31% |
| Hnrnpa2b1 | 66 | ENSMUSG00000004980 | ENSG00000122566 | HNRNPA2B1 | 5045 | 1.29% |
| Zfand5 | 15 | ENSMUSG00000024750 | ENSG00000107372 | ZFAND5 | 1153 | 1.28% |
| Eif4a2 | 33 | ENSMUSG00000022884 | ENSG00000156976 | EIF4A2 | 2537 | 1.28% |
| Pabpc1 | 15 | ENSMUSG00000022283 | ENSG00000070756 | PABPC1 | 1194 | 1.24% |
| Fyn | 11 | ENSMUSG00000019843 | ENSG00000010810 | FYN | 917 | 1.19% |
| Msl1 | 13 | ENSMUSG00000052915 | ENSG00000188895 | MSL1 | 1156 | 1.11% |
| Usp34 | 13 | ENSMUSG00000056342 | ENSG00000115464 | USP34 | 1218 | 1.06% |
| Setd5 | 10 | ENSMUSG00000034269 | ENSG00000168137 | SETD5 | 952 | 1.04% |
| Rbm5 | 15 | ENSMUSG00000032580 | ENSG00000003756 | RBM5 | 1454 | 1.02% |
| Rbm39 | 16 | ENSMUSG00000027620 | ENSG00000131051 | RBM39 | 1657 | 0.96% |
| Eif4g2 | 76 | ENSMUSG00000005610 | ENSG00000110321 | EIF4G2 | 7937 | 0.95% |
| Srsf5 | 12 | ENSMUSG00000021134 | ENSG00000100650 | SRSF5 | 1877 | 0.64% |
| Hnrnpa1 | 24 | ENSMUSG00000046434 | ENSG00000135486 | HNRNPA1 | 4579 | 0.52% |
| Ddx17 | 20 | ENSMUSG00000055065 | ENSG00000100201 | DDX17 | 4597 | 0.43% |
| Marcks | 10 | ENSMUSG00000069662 | ENSG00000277443 | MARCKS | 3508 | 0.28% |
| Ubc | 10 | ENSMUSG00000008348 | ENSG00000150991 | UBC | 26107 | 0.04% |
